# Structural visualization of septum formation in *Staphylococcus warneri* using atomic force microscopy

**DOI:** 10.1101/2020.05.24.113035

**Authors:** Hai-Nan Su, Kang Li, Xiao-Xue Yuan, Meng-Yao Zhang, Si-Min Liu, Xiu-Lan Chen, Lu-Ning Liu, Yu-Zhong Zhang

## Abstract

Cell division of *Staphylococcus* adopts a “popping” mechanism that mediates extremely rapid separation of the septum. Elucidating the structure of the septum is crucial for understanding this exceptional bacterial cell division mechanism. Here, the septum structure of *Staphylococcus warneri* is extensively characterized using high-speed time-lapse confocal microscopy, atomic force microscopy, and electron microscopy. The cells of *S. warneri* divide in a fast “popping” manner on a millisecond timescale. Our results show that the septum is composed of two separable layers, providing a structural basis for the ultrafast daughter cell separation. The septum is formed progressively toward the center with non-uniform thickness of the septal disk in radial directions. The peptidoglycan on the inner surface of double-layered septa is organized into concentric rings, which are generated along with septum formation. Moreover, this study signifies the importance of new septum formation in initiating new cell cycles. This work unravels the structural basis underlying the “popping” mechanism that drives *Staphylococcus* cell division and reveals a generic structure of the bacterial cell.

**IMPORTRANCE:** This work shows that the septum of *Staphylococcus warneri* is composed of two layers and the peptidoglycan on the inner surface of the double-layered septum is organized into concentric rings. Moreover, new cell cycles of *Staphylococcus* could be initiated before the previous cell cycle is complete. This work advances our knowledge about a basic structure of bacterial cell and provides the double layered structural information of septum for the bacterium that divide with the “popping” mechanism.

## Introduction

The septum is one of the key cellular structures during the cell division of Gram-positive bacteria. A mother cell propagates by septum formation and subsequent septum splitting (1, 2). The septum structure and formation are closely related to the means of daughter cell separation during cell division.

Gram-positive organisms have evolved various fashions for daughter cell separation. For example, division of *Bacillus subtilis* cells is mediated by the well-known gradual enzymatic splitting on the septum (3–5). Unlike this slow splitting process, cell division of *Staphylococcus aureus*, an aggressive pathogen and model coccoid Gram-positive bacterium, occurs as an ultrafast “popping” on the septum, usually on a millisecond timescale (6, 7). The fast mechanical propagation has been found not only in *Staphylococcus* and its close relative genus *Macrococcus* but also in *Actinobacteria* with diverse shapes ranging from coccoid to rod (5).

Septum separation represents a key step of bacterial cell division. Understanding the generic structure and biosynthesis process of the septum is of paramount importance for enlightening the mechanism of bacterial division. In contrast to the one-layered septum in most gram-positive bacteria, such as *Bacillus*, the septum structure of *Staphylococcus* seemed more complicated. Transmission electron microscopy (TEM) revealed a thin densely stained line located in the middle of the septum, and this structure was named the splitting system (8).. Further cryo-electron microscopy (cryo-EM) studies have led to advanced knowledge about the structure and formation of the bacterial septum (9, 10). Cryo-EM of *Staphylococcus* cells showed a low-density region sandwiched between two high-density regions in the septum, suggesting that the septum of *Staphylococcus* was composed of two layers (9). Results of scanning electron microscopy (SEM) indicated that during the separation of daughter cells, perforations on the peripheral rings of septa initiated the breakage of the peripheral rings of the septum, and eventually led to the separation of daughter cells through the ultrafast popping mechanism (7).

Advanced knowledge of septum formation in *Staphylococcus* requires extensive studies on the three-dimension structure of the whole septum, like research on isolated septa in *B. subtilis* (11, 12). In addition to the electron microscopic studies, atomic force microscopy (AFM) has been exploited in characterizing the overall structure of isolated septa from *Staphylococcus* (13) and perforations on the peripheral rings of septa (14). The peripheral rings of septa were further demonstrated to be mechanically softer than adjacent cell walls by AFM indentation (15). Despite the previous findings, the dynamic structures of the septa during septum formation and the relationship between the septum structure and cell wall synthesis remain unclear. In this study, we conducted in-depth structural visualization of the septa isolated from *Staphylococcus warneri* using AFM, which is a powerful tool in delineating the structures and spatial organizations of cell wall and biological membranes with the high resolution (16–20), combined with high-speed time-lapse confocal microscopy and electron microscopy. Comprehensive analysis of the structure and formation of bacterial septa advances our knowledge about the generic structure of the bacterial cell and the structural basis of the popping cell division.

## Results

### *S. warneri* divides in a fast popping way within milliseconds

*S. warneri* is a close relative of *S. aureus* and a coagulase-negative *Staphylococcus* species (21, 22). To examine whether *S. warneri* cells divide in the same ultrafast popping manner as *S. aureus* cells, we visualized the division process of *S. warneri* using high-speed time-lapse Andor Dragonfly confocal microscopy at 2.5-millisecond intervals following previously reported procedures (5–7). A rapid increase in cell volume occurred within 2.5 milliseconds (Fig. 1A-D), which is a characteristic feature of the “popping” division resulted from the ultrafast separation of daughter cells (5–7). Moreover, SEM images exhibit the failures in the peripheral ring (Fig. 1E) and hinges between daughter cells (Fig. 1F), the morphological characteristic of the “popping” cell division found in *S. aureus* (7). Our results demonstrated that *S. warneri* undergoes a fast “popping” daughter cell separation, on a millisecond timescale.

**Fig. 1.**
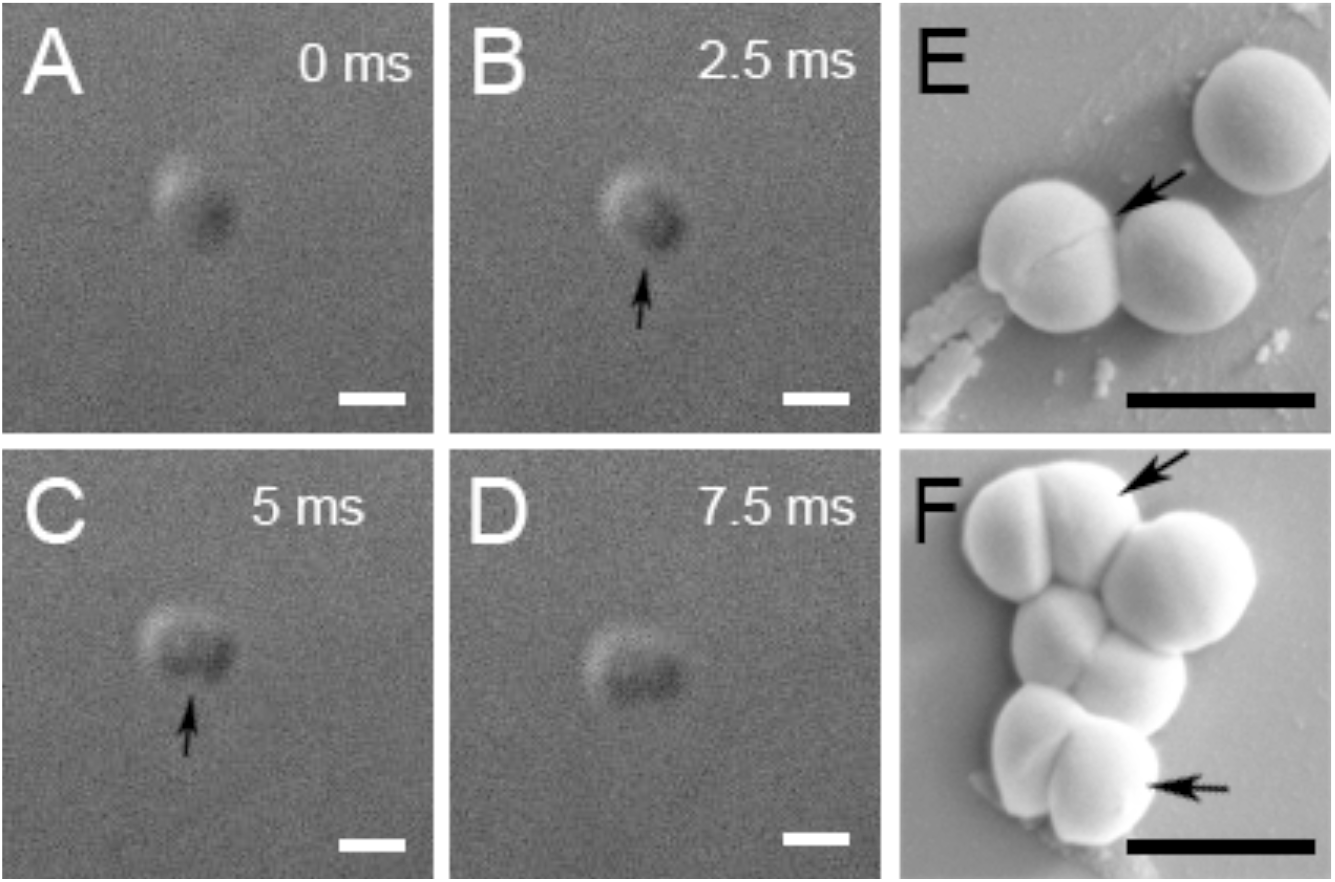
Division of *S. warneri* cells. A-D, time-lapse microscopy for daughter cell separation in *S. warneri*. Time interval for each frame was 2.5 ms. Arrows in B and C indicate the division event. Scale bar, 1 μm. E and F, SEM observations of *S. warneri* cells. The arrow in panel E indicates the mechanical failure in the peripheral ring. Arrows in F indicate the separated daughter cells connected with hinges. Scale bar, 1 μm.

### The septum is formed progressively toward the center with non-uniform thickness

To explore the structural basis that mediates the popping division, we characterized the overall structure of septa isolated from the broken sacculi fragments of *S. warneri* using AFM (Fig. S1). The septum comprises a peripheral ring and a septal disk (Fig. 2). The prepared septa contain the complete septa and incomplete septa with central gaps that vary in size, representing distinct stages of the septum formation. These results suggest that the septal disk is formed progressively toward the center until the center gap is sealed, which is similar to the synthesis process in *Bacillus* (11, 12). Analysis of the whole septa showed that the width of the annulus in the incomplete septa in every radial direction appears even (Fig. 2), indicating a simultaneous formation in all directions on the septal disk.

**Fig. 2.**
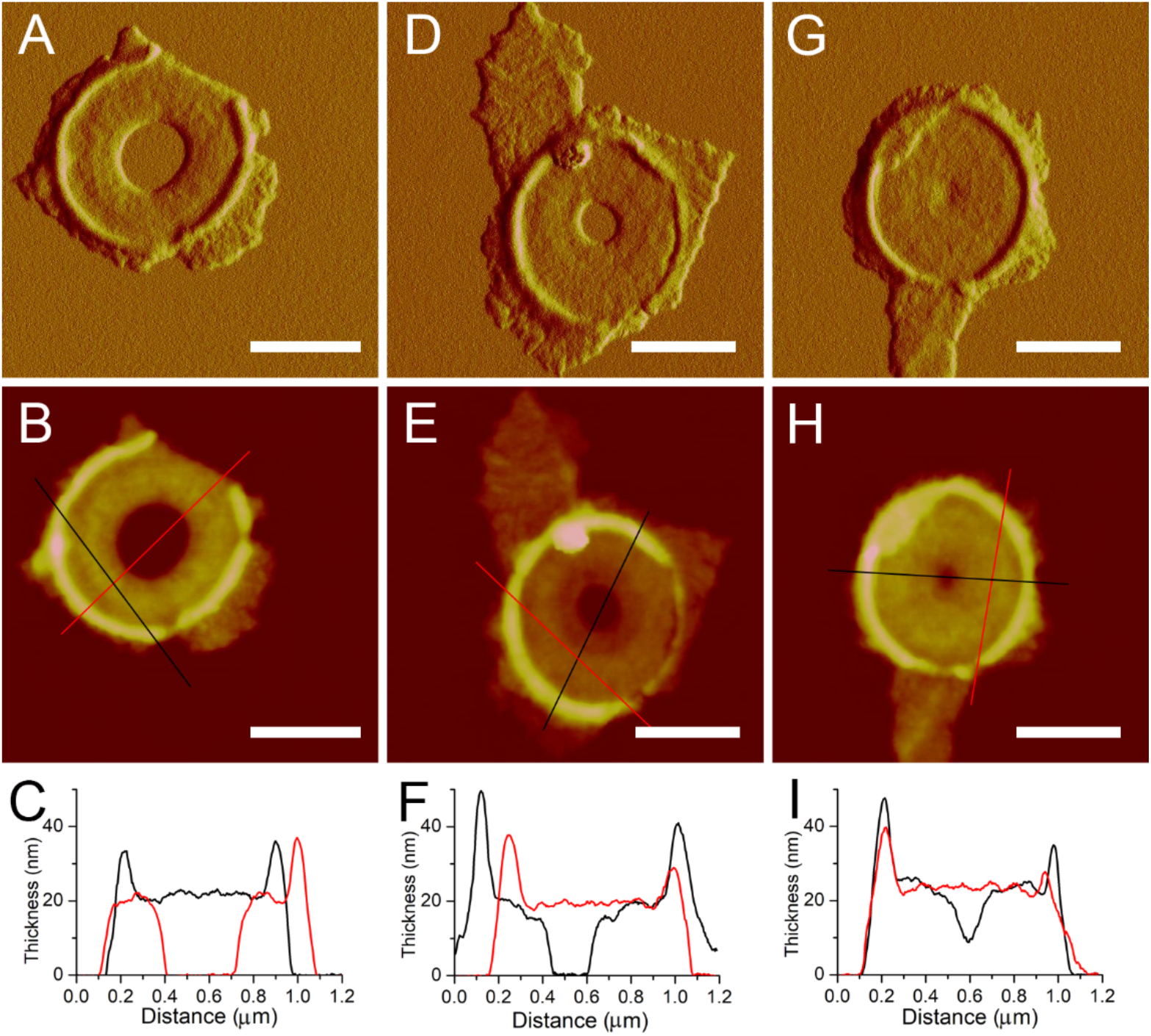
Progression of septa formation observed with AFM. First row (A, D, G), peakforce error images. Second row (B, E, H), height images. Third row (C, F, I), cross-sections in their corresponding AFM height images above (black and red lines). Scale bar, 0.5 μm.

The thickness of the incomplete septum varies across the septal disk in radial directions, with a thinner leading edge than the lagging edge, in agreement with previous observations (8, 9). The thickness distribution of incomplete septum is different from the thickness distribution in *Bacillus* (12). By contrast, the thickness of the completed septa appears relatively identical, though a thin central region was occasionally seen in the septal disk, likely representing the newly completed septum (Fig. 2G-H). Such thickness distribution suggests that the septum at the leading edge is first synthesized as a thin form; during the progress of septum formation, new cell wall materials are integrated across the developing septal surface to generate a thicker septal disk until the completed septum is formed. This dynamic synthesis process proposed based on AFM observations supported the hypothesis of septum formation suggested by thin-sections TEM of *S. aureus* cells (23).

### The septum of *S. warneri* comprises two structural layers

To elucidate the septal structure, broken septa were isolated and imaged by AFM. A total of 43 broken complete septa were found and they all showed explicitly a double-layered structure (Fig. 3A-O). The upper layers of the broken complete septa were partially lost, exposing the bottom layers. The thickness of each layer (~10 nm) is about half of the thickness of a septum (~20 nm), demonstrating that the septum is composed of two separable layers. The formation of double-layered septa could function as a structural basis for the “popping” daughter cell separation in *Staphylococcus*.

**Fig. 3.**
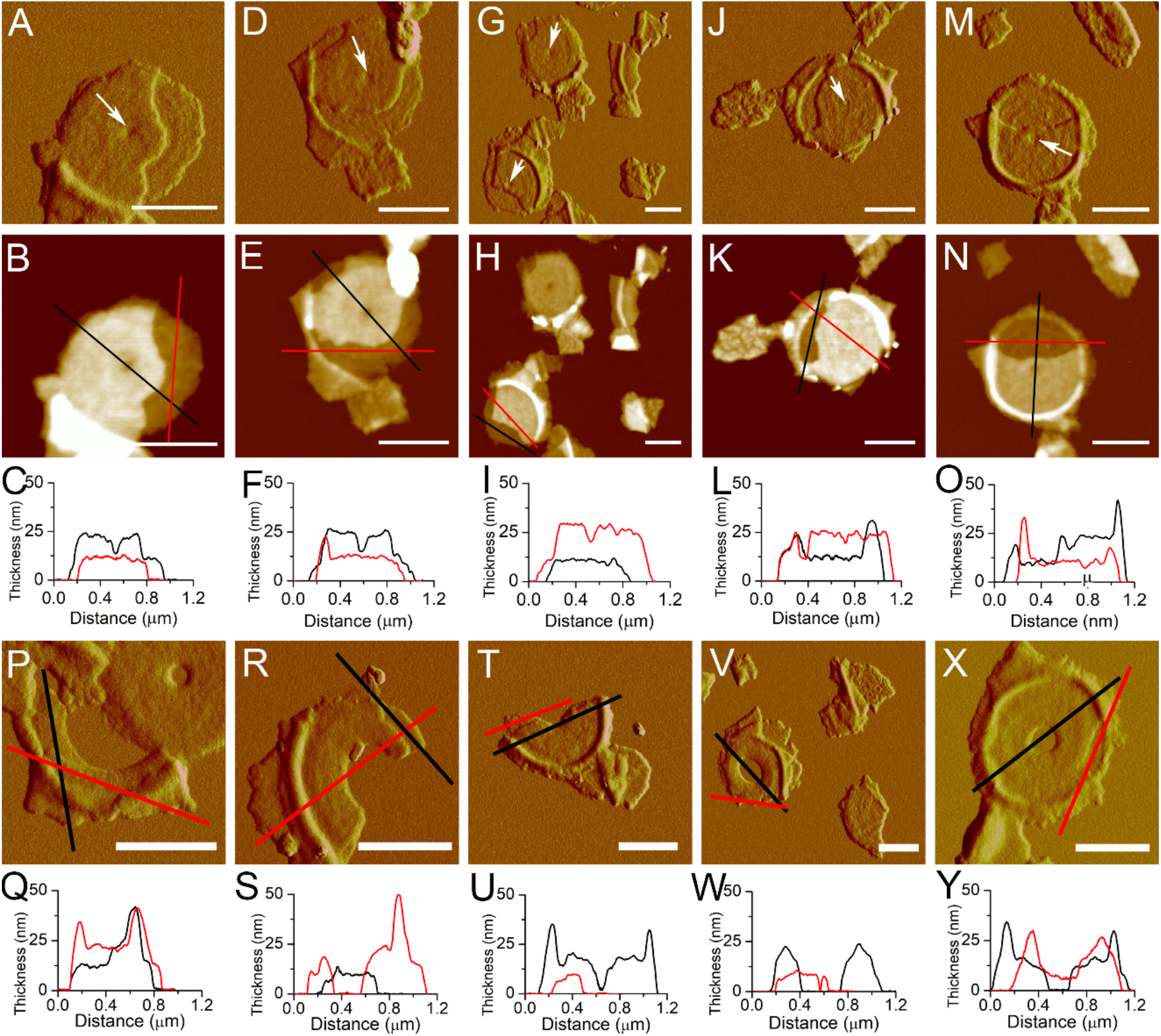
Broken complete and incomplete septa showed a double-layered structure. First row (A, D, G, J, M), peakforce error images of complete septa. Arrows indicated the central depressions. Second row (B, E, H, K, N), height images of complete septa. Third row (C, F, I, L, O), cross-section analysis of complete septa in their AFM height images above. Fourth row (P, R, T, V, X), peakforce error images of incomplete septa. Fifth row (Q, S, U, W, Y), cross-section analysis of incomplete septa in their corresponding AFM height images above. Positions for the cross-section analysis were indicated by black and red lines. Scale bar, 0.5 μm.

AFM imaging of broken completed septa could not allow us to answer whether (i) the double-layered structure was formed during the formation of the septum, or (ii) a completed single-layered septum was first formed and then split into two layers. To address these questions, we carried out AFM analysis on the broken incomplete septa at various stages of septal formation. The double-layered structures were visible from the leading edge to the lagging edge of the incomplete septa (Fig. 3P-Y), suggesting that the double-layered structure was formed during septum formation.

### Concentric ring structures on new cell walls were formed prior to separation of the septum

We identified the concentric rings as the characteristic structures of new cell walls in *S. warneri* cells (Fig. S2), in line with other *Staphylococcus* species in previous studies (14, 24). Similar structures have also been discerned in the cell walls of other coccid Gram-positives (25–27). Then, we imaged the isolated new cell wall fragments of *S. warneri* using AFM. Of the 117 isolated new cell wall fragments characterized, which have a flat disc shape with the thickness of ~10 nm (Fig. S3), 65 fragments (56%) showed concentric ring structures on the cell wall surfaces (Fig. 4A-B), whereas the surface features of the other fragments 44% were comparatively less clear (Fig. 4C-D). It is likely that the featured surface and relatively featureless surface represent the two distinct oriented sides of new cell walls. High-resolution AFM images showed that the ring structures were tightly associated, with an average bandwidth of ~16 nm (*n* = 30) (Fig. 4E-H).

**Fig. 4.**
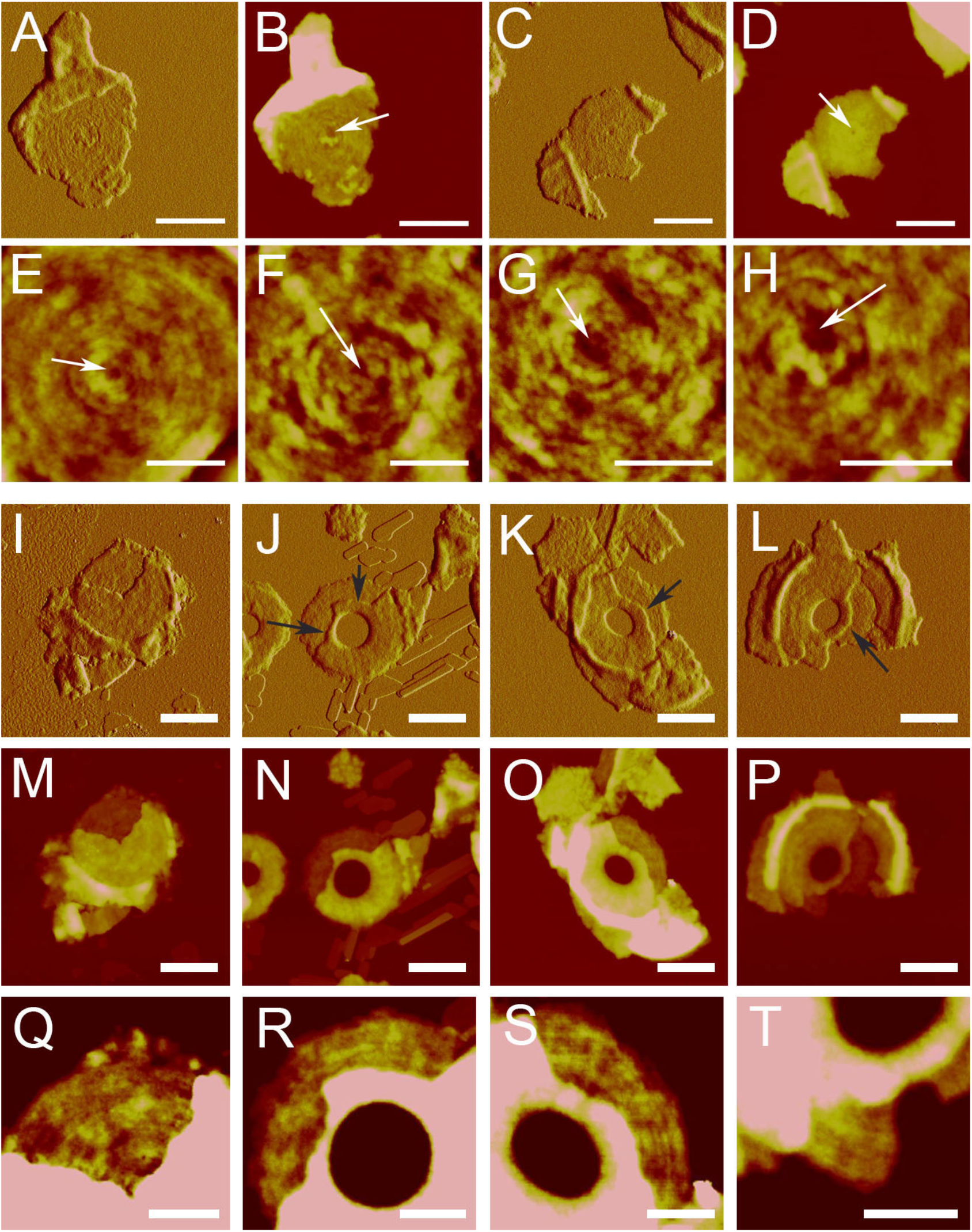
Concentric ringed structures on the surface of new cell walls and the inner surface of double-layered septa. First row, AFM images of isolated new cell walls. A and B, new cell wall with concentric rings. Scale bar, 0.5 μm. C and D, new cell wall with unclear surface features. Scale bar, 0.5 μm. A and C are peakforce error images; B and D are height images. Second row, high-resolution AFM height images of the concentric rings on new cell walls. Scale bar, 200 nm. White arrows indicated the central depressions. Third row, peakforce error images of broken septa. Black arrows indicated fractures. Scale bar, 0.5 μm. Fourth row, height images of the broken septa in the corresponding peakforce error images above. Scale bar, 0.5 μm. Fifth row, zoomed-in images from the corresponding height images above. Scale bar, 200 nm.

When the concentric structures in new cell walls were generated remains unclear. It is known that new cell walls were formed by separation of the double-layered septum. This information was further confirmed by the fact that about half of the isolated new cell wall fragments showed concentric ring structures while the other half did not (Fig. 4). No concentric rings were determined on the outer surface of the septum (Fig. 2). However, in the broken complete septa, concentric ring structures appeared in the exposed inner surface (Fig. 4I, Fig. 4M, and Fig. 4Q), indicating that each layer of the septum had one relatively featureless side and one featured side with concentric rings.

We next questioned whether the concentric rings were formed after completion of the septum formation or along with the septum formation. AFM images of the exposed inner surface of the incomplete septa reveal the presence of concentric rings on the inner surface of the double-layered incomplete septum (Fig. 4J-L, Fig. 4N-P, Fig. 4R-T). The results demonstrated that the concentric ring structure was formed during septum formation.

Visualization of the concentric rings provides clues of how peptidoglycan is organized in the septum. The concentric ring organization of peptidoglycan could be generated with the progressive formation of the septum toward the center of the septal disk (Fig. 2). Moreover, we found that fractures on the broken septa layers often appear along the direction of the concentric rings (Fig. 4J-L, black arrows), further suggesting that the concentric rings might not only be a surface feature but also reflect the intrinsic peptidoglycan organization in cell walls.

### A connection between two layers of the incomplete septum at the interior leading edge

Among the broken incomplete septa that we characterized (*n* = 30), partial upper layers were absent and bottom layers were exposed. In most cases (29 out of 30), the broken incomplete septa showed no sign of separation of two layers at the interior edge. In some cases, the fractures could be were quite close to the interior edge, as depicted in Fig. 4J-L (black arrow). However, the interior edge remained intact. It allowed us to assume that the two layers were separable but separation at the interior leading edge may be hard, implying a strong connection between the two layers of the incomplete septum at the interior edge, consisting with previous cryo-EM observations (9).

To further verify this hypothesis, we investigated the incomplete septa that have lost their peripheral rings. The peripheral ring is the only site that could form a structural connection between septum layers, as shown in previous SEM images (7) and cryo-EM images (9) of the dividing *Staphylococcus* cells. When the peripheral ring is removed, connections between two layers at the peripheral sites should be lost. Therefore, since the two layers of the septum could be separated as proved by our AFM observations, for the incomplete septa without peripheral rings, it is expected that two layers could be completely separated apart if there are no connections between them at the interior edge, and *vice versa*. We found that in total 17 incomplete septa whose peripheral rings were completely lost, no separation of the two layers of the incomplete septa was observed (Fig. 5A-D, Fig. S4). Moreover, no separated single layers of incomplete septa were detected during our work.

**Fig. 5.**
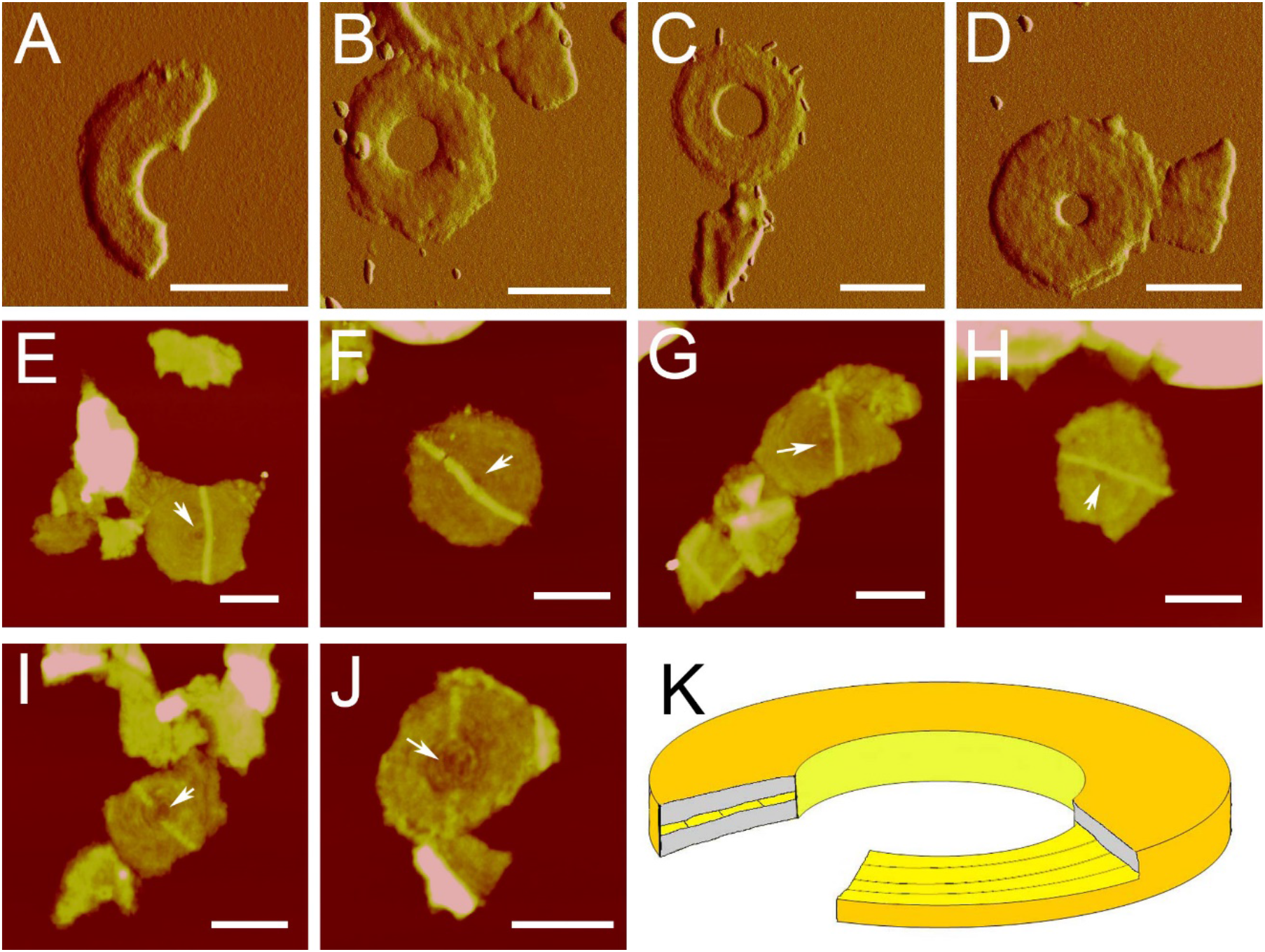
Analysis of the connection at the interior leading edge of incomplete septa. First row, peakforce error images of incomplete septa that had lost peripheral rings. Scale bar, 0.5 μm. Second row, height images of new cell walls with complete belts. Scale bar, 0.5 μm. I and J were height images of new cell walls with discontinuous belts. Scale bar, 0.5 μm. White arrows indicated the central depressions. K, a structural model of an incomplete septum.

In *Staphylococcus*, new septa are synthesized perpendicular to the previous division plane crossing the previous new cell wall (6, 13). We visualized 48 new cell wall fragments with belt-like structures growing across the new cell walls (Fig. 5E-H), which may represent the initial synthesis of new septa (13). These belt-like structures do not exhibit a two-layered structure but are complete belts that are sealed at the top (Fig. 5E-H). This result further suggests that the two layers of incomplete septa could be connected at the interior leading edge.

### The central depression is an inherent structure in the septum and new cell wall

AFM images revealed the existence of the central depression in almost all the complete septa (Fig. 3A-O). One may question whether the central depression we observed is an inherent structure or is resulted from the incomplete synthesis of the septum. To address this question, we analyzed the structures of the new cell walls. New cell walls are transformed from the complete septa after daughter cell separation. The possibility of incomplete synthesis of septa could be excluded by analyzing the structure of new cell walls. We found that the central depression is always present in the centers of new cell walls enclosed by the concentric rings (Fig. 4A-H and Fig. 5E-H). Moreover, the depression structures have also been seen in the center of new cell walls in intact bacterial cells (14, 24). Therefore, it is unlikely that the central depression was resulted from the incomplete synthesis of the septum but may represent an intrinsic structure of the septum and new cell walls. This depression feature has not been observed in previous TEM images (28–32), as identifying this structure requires proper sectioning across the center of septa. Our AFM results provide insight into the septum structure and raise an open question of how the last step of septum synthesis is performed.

An interesting observation was the initiation of the synthesis of a new septum. When new septum synthesis was initiated, they always choose a path to avoid crossing the central depressions (Fig. 5E-H). Although the physiological function of the central depression remains unclear, the results indicate that the central depression is mechanically unfavorable for the synthesis of a new septum. It is possible that the central depression hampers proper localization and interactions of protein complexes that are contributing factors in septum synthesis. In some cases, cells tend to build a new septum in the direction across the center depression. However, synthesis was halted in the vicinity of center depressions (Fig. 5I-J). No other types of discontinuous belt structures were observed in our work. These results suggest a negative effect of the center depression on new septum synthesis.

### A new round of cell cycle could be initiated before the septum is separated

The cell cycle of *Staphylococcus* involves three phases (6). In phase 1, cells are slightly elongated without formation of the septum. Then, the cells initiate and complete the septum formation in phase 2 (Fig. 6G). In phase 3, cells with complete septa are elongated and ultimately divide into two daughter cells (6). After division, a new round of the cell cycle in *Staphylococcus* cells commences. We found that some *Staphylococcus* cells adopt different fashions of their cell cycles. 32 out of 115 completed septa have unequivocal belt-like structures (Fig. 6A-F). The thickness of these sacculi fragments is ~20 nm, indicating that they are septa instead of new cell walls. More importantly, some broken septa with belt-like structures exhibit double-layered structures (Fig. 6A-F), suggesting that these belt-like structures were synthesized on real septa. These belt-like structures were observed only on complete septa and never on incomplete septa, probably revealing that the belt structures were synthesized after the septa were complete. Formation of such a belt structure showed initiation of the synthesis of new septum before the old septum was separated. These results indicate that a new round of the cell cycle could commence before the last one is complete (Fig. 6H).

**Fig. 6.**
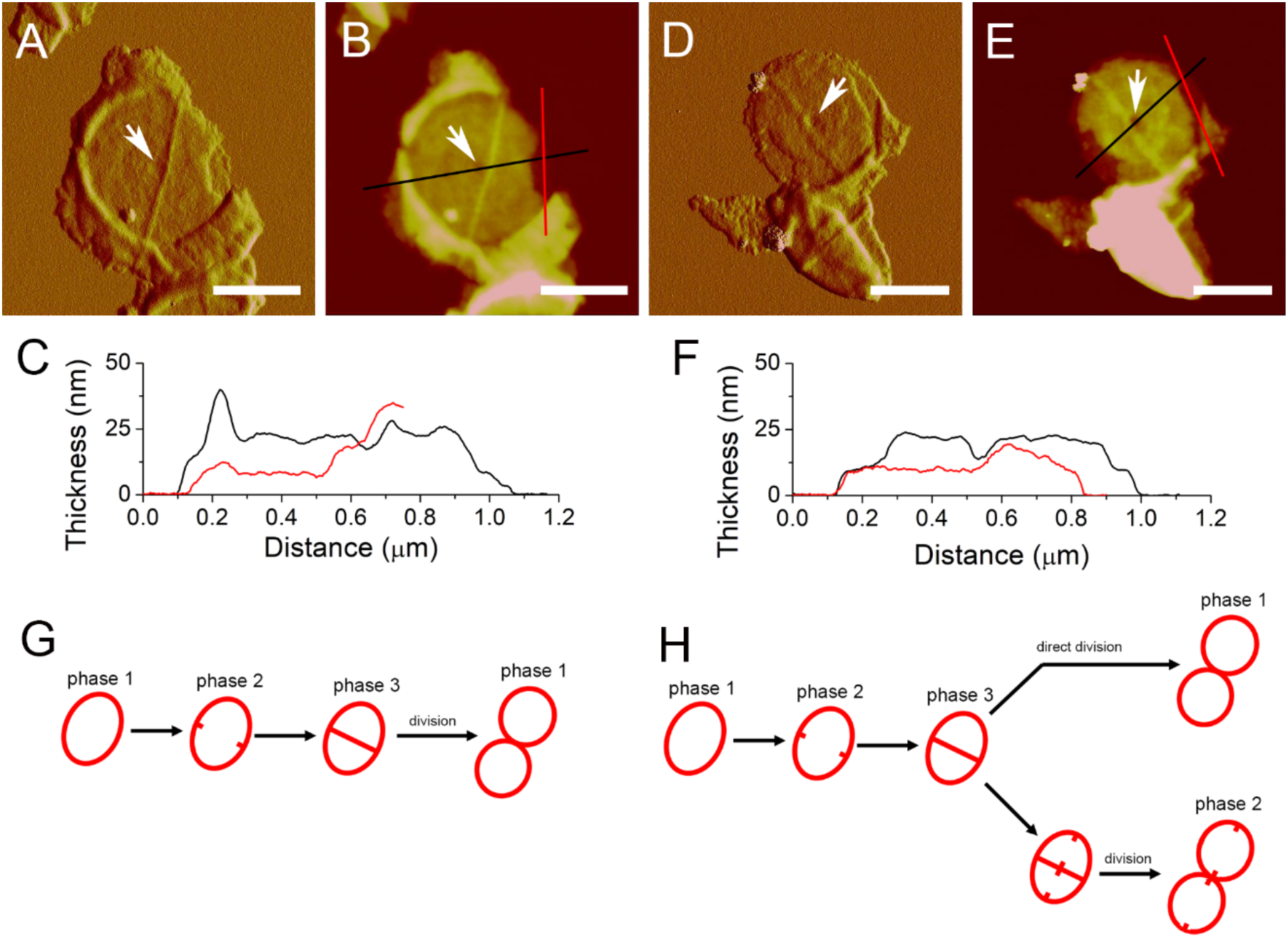
Belt structures formed on the complete septa. A and D were peakforce error images, while B and E were height images. C and F were section analyses on B and E, respectively. Positions for section analyses were indicated with black and red lines. White arrows indicated the central depressions. Scale bar, 0.5 μm. G, the previous model of the cell cycle in *Staphylococcus*, adopted from Monteiro et al (6). H, a new model of the cell cycle in *Staphylococcus* based on this work.

## Discussion

The septum is a key cell wall structure that is of physiological significance for cell division of Grampositive bacteria. Here we conducted extensive characterization of the dynamic structures of septa from *S. warneri*, which divides in a fast “popping” mechanism on a millisecond timescale. A striking feature is that the septum is composed of two separable layers (Fig. 5K), which may lay the mechanistic foundation for bacteria like *Staphylococcus* to undergo daughter cell separation in an extremely fast “popping” fashion (6, 7).

The peptidoglycan on the newly synthesized cell wall of *Staphylococcus* is organized to form concentric ring structures, in agreement with the previous finding (14, 24). Our results suggested that such structures in new cell walls may be formed before the septum is separated. We demonstrated that the concentric ring structures appear in the inner surface of the septal peptidoglycan and could be formed with the progressive formation of the septum toward the center of the septal disk. The concentric ring structures in septal peptidoglycan resemble the so-called “splitting system” observed with thin-section TEM of bacterial cells (8). The nature of the splitting system has remained elusive over the past decades. This work unraveled that the peptidoglycan is organized as the concentric ring structures in the septum.

Bacterial cell cycle requires several complex cellular processes that should be tightly regulated and precisely coordinated (2, 33, 34). Formation of the septum is an important process in the bacterial division and should occur in time to ensure the quality of propagation (33). We found that part of bacterial cells initiated the next round of the cell cycle in advance before the previous one is complete. It remains unknown whether such an unusual cell cycle was resulted from the failure of regulation, or it is an adopted strategy in response to internal or environmental variations. This work indicated that the arrangement of a cell cycle in *Staphylococcus* differs greatly at the single-cell level. Elucidating how bacterial cells enter the next round of the cell cycle is of fundamental importance for a deeper understanding of the regulatory mechanism underlying bacterial growth and division.

## Materials and Methods

### Purification of sacculi

Sacculi were purified as described previously (11, 13). Briefly, *S. warneri* was grown in Luria-Bertani (LB) medium at 25°C overnight with agitation. Cells were collected and boiled for 7 min. For isolating septa or broken sacculi, cells were broken by ultrasonication (200~400w). Then cells were treated with boiling in SDS (5% w/v), RNase (0.5 mg/ml), DNase (0.5 mg/ml) and pronase (2 mg/ml). Removal of accessory polymers was achieved by incubation in 48% v/v HF at 4°C. Purified sacculi were washed with MilliQ water at room temperature.

### AFM

AFM imaging was carried out by using a Multimode VIII AFM with Nanoscope V controller (Bruker AXS, Germany). AFM imaging in air condition was carried out in Scanasyst mode. Silicon nitride cantilevers (XSC11/AL BS, NanoAndMore Corp, USA) were used for imaging in ambient conditions.

### SEM

Samples were fixed in glutaraldehyde and osmium tetroxide, and then dehydrated by a series of increasing concentrations of ethanol (10%, 30%, 50%, 75%, 90%, 95%, and 100%). Then samples were dried with carbon dioxide in a critical point dryer EM CPD 300 (Leica, Germany), and sputter-coated with gold-platinum in a Sputter Coater 108 (Cressington, UK) for 240 seconds and examined with a Quanta FEG 250 (FEI, USA) at 10 kV.

### High-speed time-lapse confocal microscopy

Two-dimensional (2D) time-lapse imaging was performed on Andor Dragonfly confocal system. The microscope was Leica DMi8 with 100× (NA 1.4) oil-immersion objective. Cells were imaged at room temperature. The camera was Andor Zyla 4.2 Plus USB version, and capture speed was 400 fps at 512 × 512 pixel array size. Camera pixel size was 6.5 μm.

## Acknowledgements

We would like to thank Haiyan Yu, Xiaomin Zhao, Sen Wang and Qi Chen from State Key Laboratory of Microbial Technology of Shandong University for technical help in this work. This work was supported by the National Science Foundation of China (31900023, 31570066), National Key R&D Program of China (2018YFC1406701), the Program of Shandong Taishan Scholars (TS20090803), and Young Scholars Program of Shandong University (2017WLJH22), UK Royal Society (UF120411, URF\R\ 180030) and Biotechnology and Biological Sciences Research Council Grants (BB/M024202/1, BB/R003890/1). The funders had no role in study design, data collection and analysis, decision to publish or preparation of the manuscript.

## Author contributions

HNS and LNL designed the research; HNS, LNL, KL, XXY, MYZ and SML performed the research; HNS, XLC and LNL analyzed the data; HNS, YZZ and LNL wrote the paper.

## Supplementary Information

**Fig S1.**
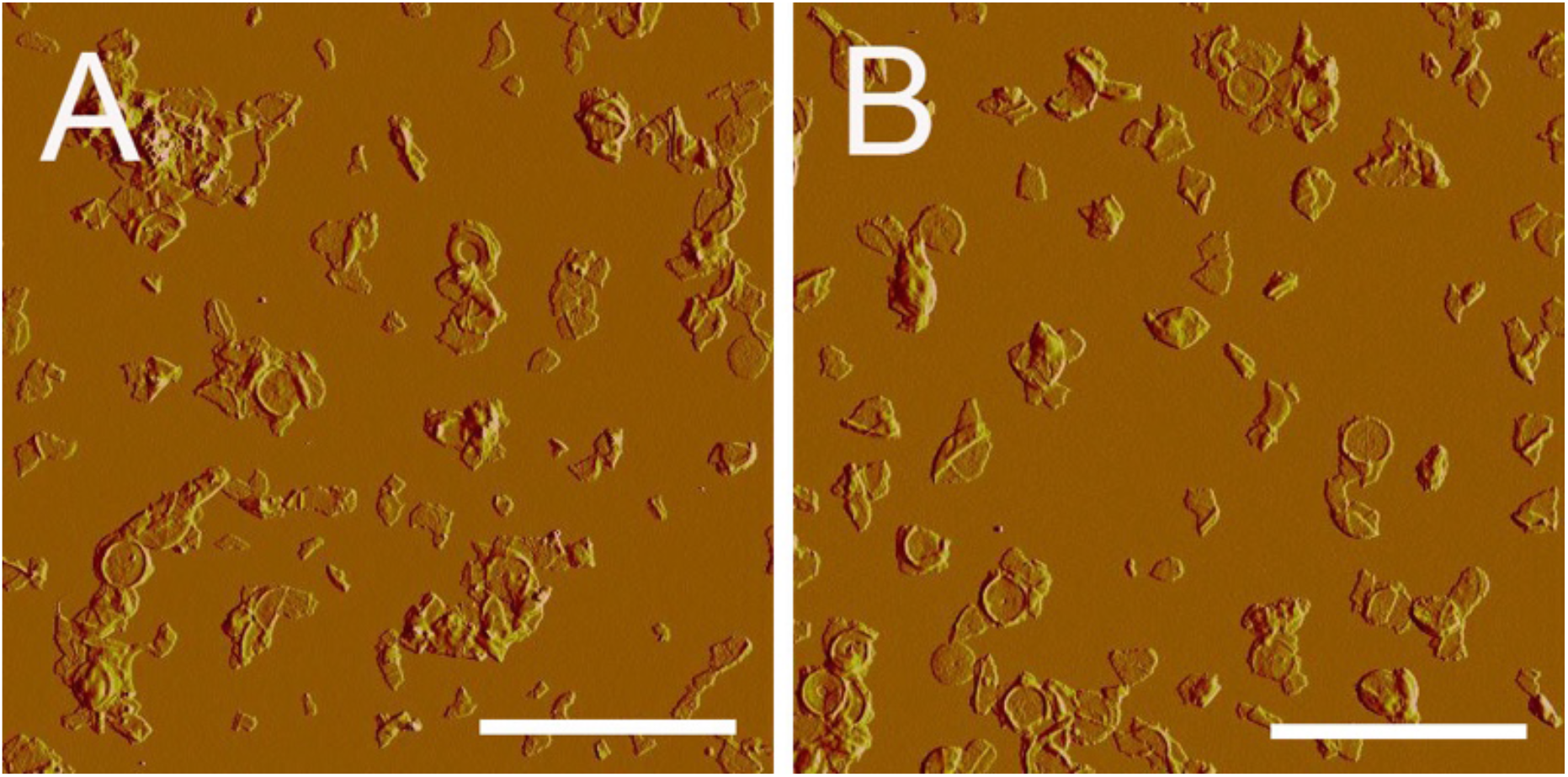
Isolated sacculi fragments from *S. warneri*. A, B, AFM peakforce error images of the overview of sacculi fragments. Scale bar, 5 μm.

**Fig S2.**
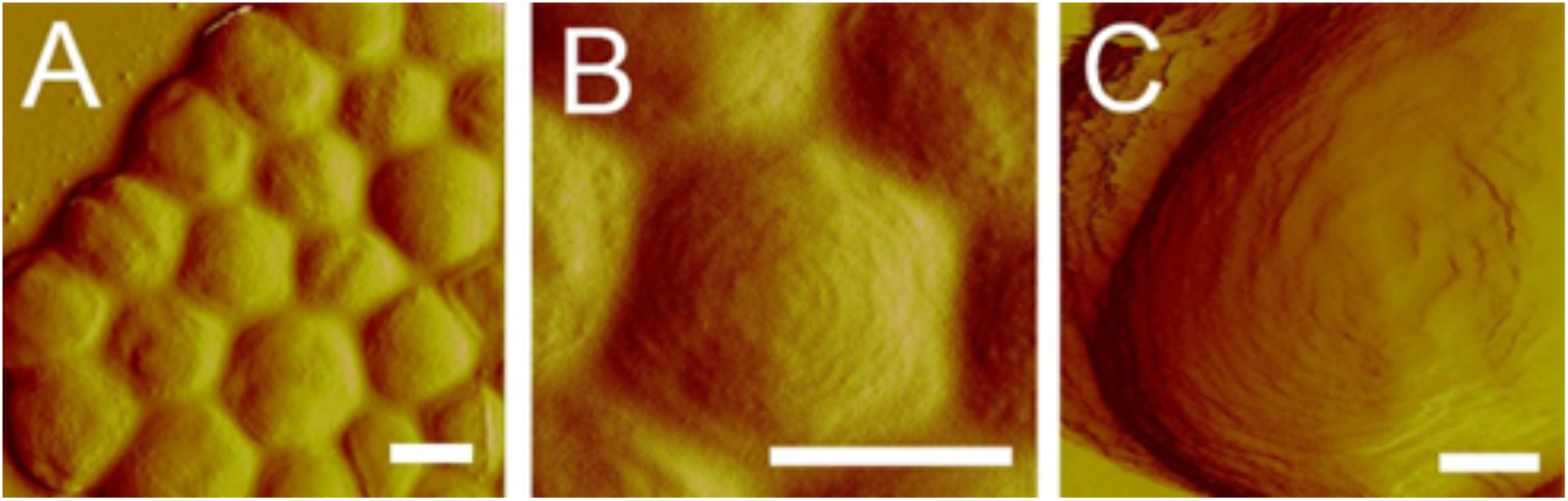
Concentric ring structures on the surface of *S. warneri* cells. A and B, *S. warneri* cells observed in air condition. C, *S. warneri* cell observed in liquid condition. AFM imaging in liquid condition was carried out in contact mode, using Multimode VIII AFM with Nanoscope V controller (Bruker AXS, Germany) with SNL-10 cantilevers (Bruker, Germany). All images were peakforce error images. Scale bar, 0.5 μm in A and B and 200 nm in C.

**Fig S3.**
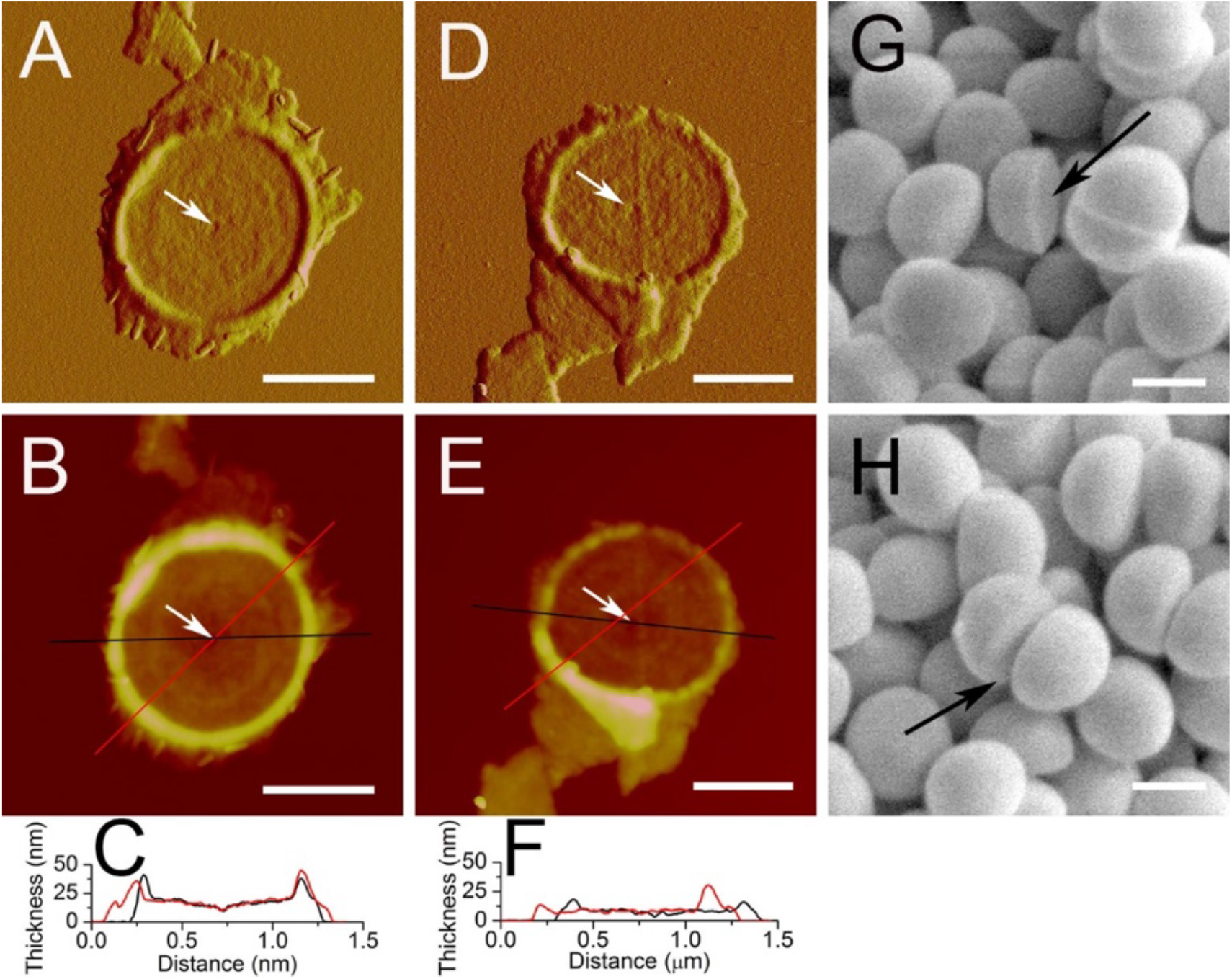
Flat disc-shaped sacculi fragments from *S. warneri*. A and B, septa with a thickness of about ~20 nm. Scale bar, 0.5 μm. D and E, new cell wall with a thickness of about ~10 nm. A and D were peakforce error images. B and E were height images. Scale bar, 0.5 μm. C and F are section analyses in B and E, respectively. Section positions are indicated by lines. White arrows indicate the central depressions. G and H, SEM images of *S. warneri* cells. Scale bar, 0.5 μm. Black arrows indicate flat new cell wall surfaces. When new cell walls were just formed from split septum in a recent division, and yet to be reshaped into hemisphere, they exhibited a flat disc-like structure, which could be confirmed from scanning electron microscopy images of *S. warneri* cells.

**Fig S4.**
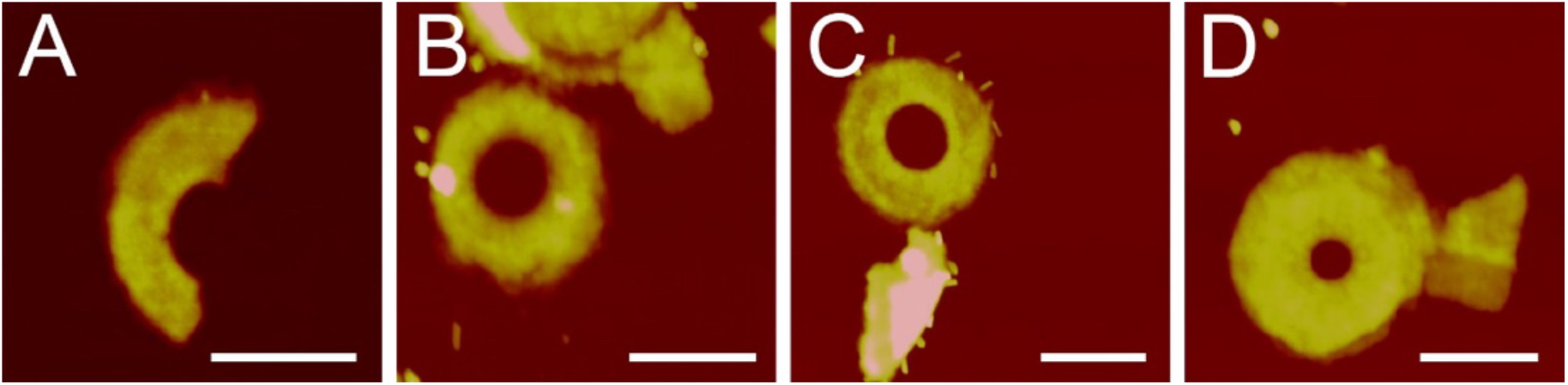
Incomplete septa that had lost peripheral rings. AFM height images of septa, corresponding directly to the peakforce error images in Fig. 5A-D. Scale bar, 0.5 μm.

